# A high-cholesterol diet leads to faster induction of general anesthesia in two model animals: *D. magna and C. elegans*

**DOI:** 10.1101/2022.11.30.518590

**Authors:** K. Carlo Martín Robledo-Sánchez, J. C. Ruiz-Suárez

## Abstract

General anesthesia (GA) has been under scientific scrutiny since its discovery more than a century ago, resulting in conceptually different proposed mechanisms to explain its origin and operation. Two mechanisms stand out: the lipid and the protein hypothesis. The Meyer-Overton rule (the more anesthetics dissolve in octanol, the greater their action) backups the first hypothesis, while the ligand-receptor interaction, specifically on ion channels, sustains the second. A recent study on *Drosophila melanogaster* draws attention to the possibility that both paradigms come together to explain GA synergistically, with the important caveat that this hybrid mechanism lies in the existence of lipid rafts in which cholesterol plays an essential role. Using two model organisms, the water flea (*D. magna*) and the nematode *C. elegans*, we give a further step to clarify this puzzle by carrying out anesthetic experiments with xenon and nitrous oxide. First, the obtained dose-response curves are very steep, implying that Hill coefficients greater than one are needed to describe them correctly, supporting an unspecific action mechanism. Second, we show that the animals’ response to both gases is influenced by a cholesterol diet modification, thus proving that this lipid promotes anesthetic induction. Our findings reenforce the idea that GA is driven by an allosteric induction rather than selective actions on single-target receptors.

---

Despite being safe and reliable in the medical praxis around the world, GA is an unsolved scientific problem so the search for the correct mechanism behind its action remains a current and active effort [1–5]. For now, although the most accepted theory is that GA is based on the effects anesthetics produce on transmembrane proteins called ion channels [6], an old hesitation remains: since anesthetic agents are hydrophobic molecules, how can they efficiently couple with proteins through specific interactions? Therefore, the idea that GA obeys an action mechanism based on specific receptors has not been fully accepted [2,7-8], upholding the notion that anesthetic molecules inhibit transport currents in nerves not because they act selectively on target receptors, but because they act on unspecific targets. In this regard, a recent study on *Drosophila melanogaster* highlights the possibility that the lipid and protein paradigms could join to synergically explain GA. Indeed, it has been found by Pavel *et al*. [3] that anesthetics disrupt lipid rafts on *Drosophila melanogaster*, activating TWIK-related K^+^ channels (TREK-1), thus constituting a mechanism distinct from the usual receptor–ligand interaction to explain inhaled anesthesia. Furthermore, they also found that cholesterol depletion in the lipid rafts triggers anesthesia by the activation of TREK-1. Both events are equivalent since cholesterol depletion also disrupts the rafts. Recently, it has been reported that the anesthetic effect of Xe could be mediated by the sequestration of cholesterol [9], an event that is equivalent to depletion.

The purpose of the present article is to report our recent findings obtained in GA experiments with gaseous anesthetics employing two organisms: the crustacean *Daphnia magna* (DM) and the nematode *C. elegans* (CE). DM has a primitive nervous system that shares several similitudes with mammals, like the presence of the neurotransmitters acetylcholine, serotonin, dopamine, epinephrine [10]. A recent work also suggests that DM shares neurological mediated pathways with humans and that some of them are similarly affected by psychiatric drugs [11–14]. *C. elegans*, on the other hand, represents an invaluable model in anesthesia experiments since it shares many genes with mammals’ cells [15-16].

Although NMDA glutamate receptor is necessary for pharmacological action of xenon and nitrous oxide, we demonstrate that the nmr-1(null) CE mutant can be anesthetized using both gases. Furthermore, since proteins are generally located in specific lipid domains known as lipid rafts composed meanly by cholesterol [3, 17-19], we emphasize on the anesthetic effects as the cholesterol diet of the animals is modified. In CE, cholesterol accumulation is most prominent in the pharynx, nerve ring, excretory gland cell, gut, and germ-line cells [20]. This multipurpose molecule determines physic properties of the cell membrane as fluidity and ion permeability [21] and serves as a precursor and co-factor for other signaling molecules [22-23]. Furthermore, this lipid offers many possibilities for the regulation of the membrane-embedded, since depending on the kind of receptor, cholesterol can strongly influence the affinity state for binding and signal transduction. However, one of the most important functions of cholesterol in cell membrane is the stabilization of receptors in defined conformations related to their biological functions [24-26]. Indeed, cholesterol is an essential component of lipid bilayers since it can combine with sphingolipids and gangliosides to form specific lipid microdomains called rafts [27-29]. Lipid rafts are located on the cell surface and are implicated in protein sorting. These dynamic assemblies change their size and composition in response to intra- or extracellular stimuli [30]. Furthermore, it has been suggested that they are attractive candidates as platforms that coordinate signal transduction pathways with intracellular substrates [31].

## Results

The anesthetic effects of xenon and nitrous oxide were evaluated in DM and CE using a high-pressure cell installed under a microscope, see *SI Appendix*, Fig. S1. As shown in Fig. 1A-C, changes in motility were obtained from the autocorrelation of images obtained during the anesthetic process (see *SI Appendix*, movies S1, and S2, Figs. S2 and S3, and details in *Materials and Methods*). An autocorrelation value (ϕ) close to 0 means full movement, which represents a normal response. When this parameter approaches 1, the motility is negligible. In summary, the quicker ϕ drops from to 0, the larger the motility of the animals (Figs. 1 A-C).

**Fig. 1.**
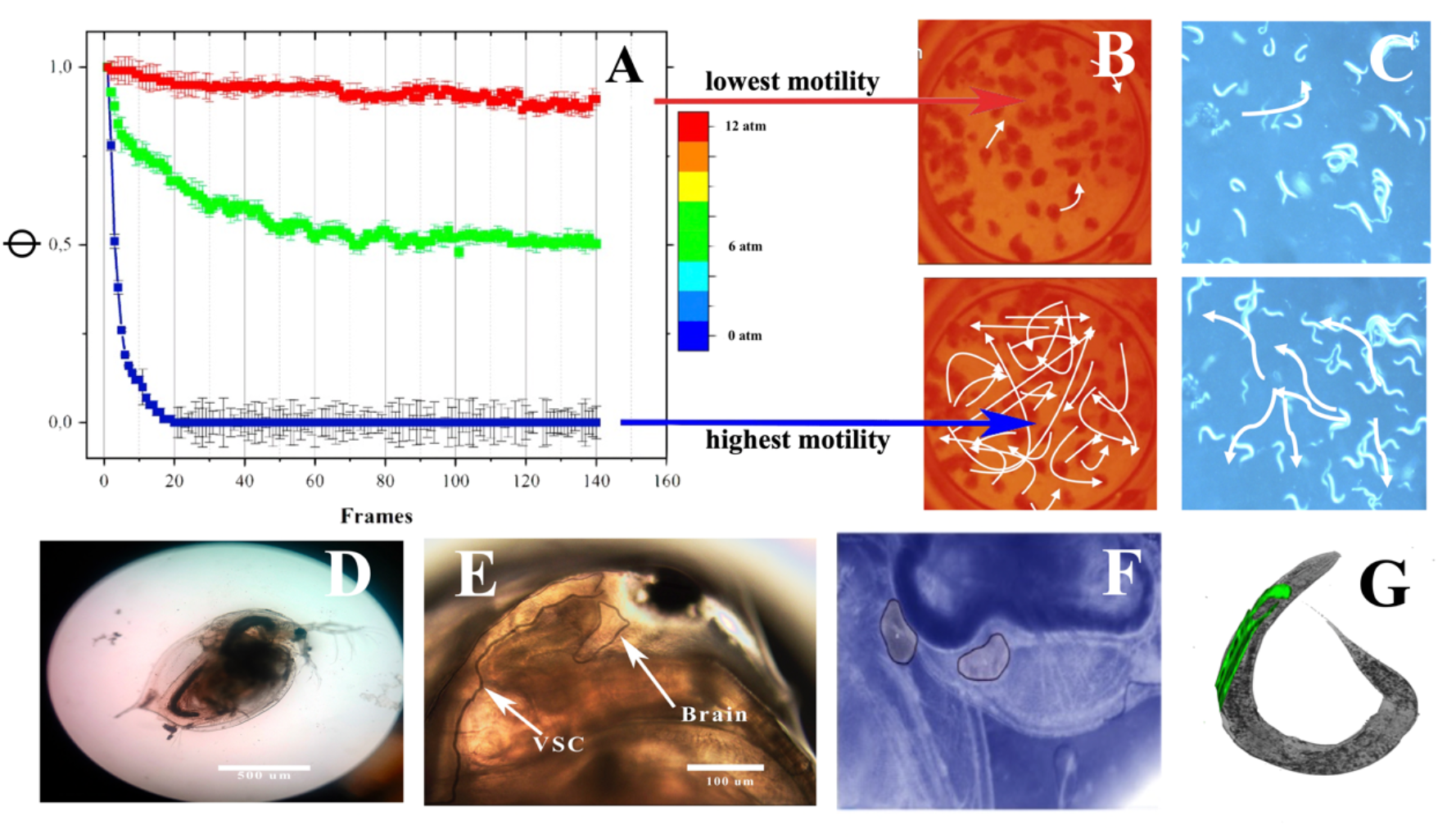
Autocorrelation curves (A) for various stages of *D. magna* (B) and *C. elegans* (C). Photograph of a single *D. magna* on the microscope (D), head (E), hearts of two contiguous fleas (F), and a single nematode (G).

Figure 2 depicts the dose-response curves for DM (A, D) and CE (B, C, E, and F), obtained by processing the autocorrelation curves using the mathematical expression given in *Materials and Methods*. The best fits of the data (solid lines, Fig. 2) were obtained using Hill’s equation [32]:

**Fig. 2.**
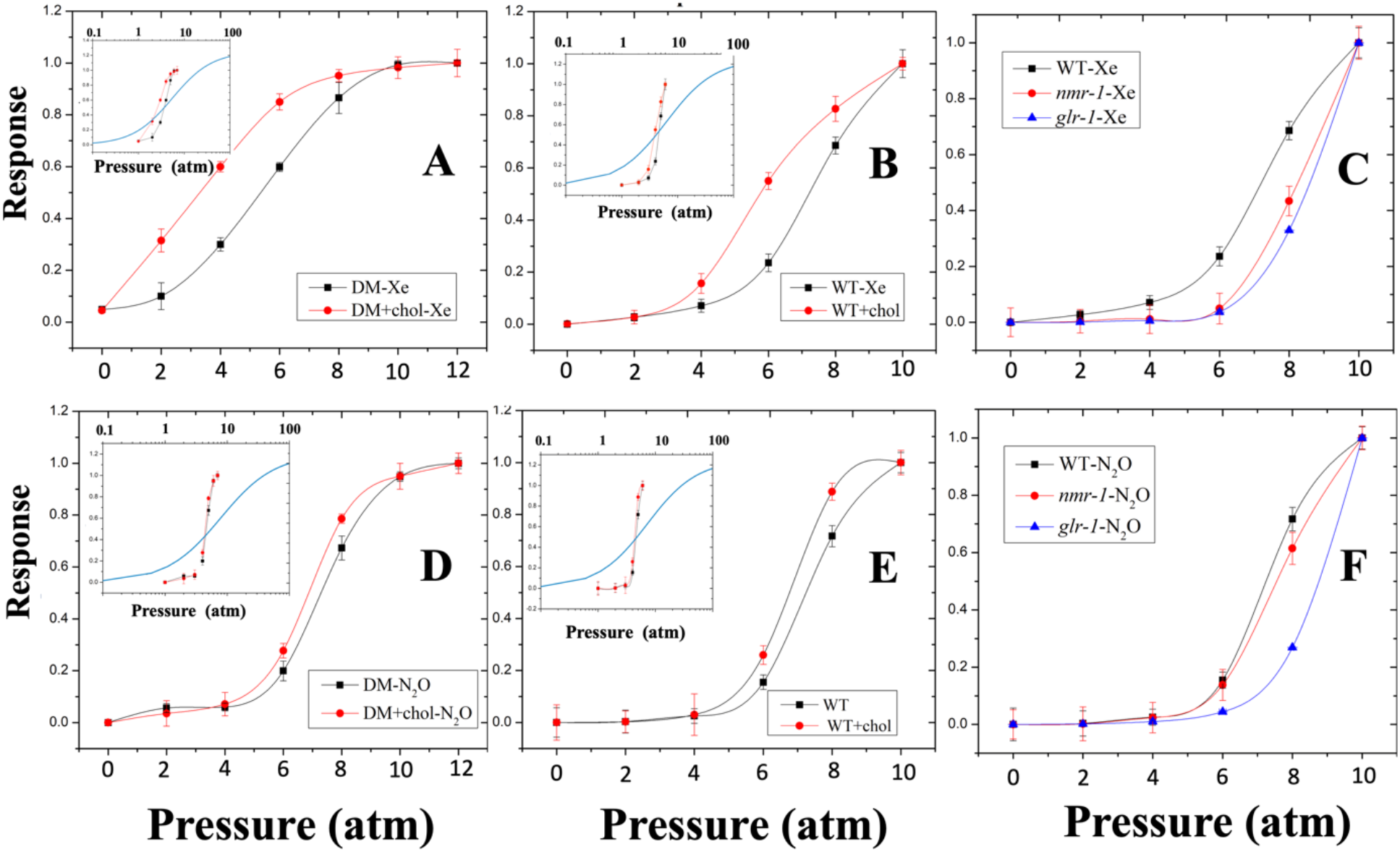
Dose-response curves obtained using Eq. 2 and autocorrelation curves displayed in *SI Appendix*, Figs. S3 and S4: where the definitions of the dose and response are detailed in *Material and Method*s. Doses-response for DM using Xe with and without cholesterol diet (A). Dose-response for CE wild type using Xe with and without cholesterol diet (B). Dose-response for CE wild type, *nmr-1* and *glr-1* using Xe (C). Dose-response for DM using N_2_0 with and without cholesterol diet (D), Dose-response for CE wild type using N_2_0 with and without cholesterol diet (E), and Dose-response for CE wild type, *nmr-1* and *glr-1* using N_2_0 (F).

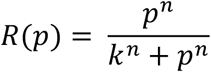

where *p* is the pressure (dose) measured in atm or bar, k is the pressure where the half-maximal effect concentration [EC]_50_ is achieved, and *n* the Hill coefficient, which determines the steepness of the logistic curve. We depict in Table 1 the values for k and n in all the cases.

**Table 1.** Parameters k and n of Hill’s equation that best fit the data in Fig. 2.

| Animal | DM | DM | DM | DM | CE wt | CE wt | CE wt | CE wt | CE vm487 | CE vm487 | CE kp4 | CE kp4 |
| --- | --- | --- | --- | --- | --- | --- | --- | --- | --- | --- | --- | --- |
| Gas | Xe | N<sub>2</sub>O | Xe-Chol | N<sub>2</sub>O-Chol | Xe | N<sub>2</sub>O | Xe-Chol | N<sub>2</sub>O-Chol | Xe | N<sub>2</sub>O | Xe | N<sub>2</sub>O |
| $k$ (atm) | 7.02 | 7.2 | 3.5 | 6.8 | 7.1 | 7.2 | 4.9 | 6.1 | 8.2 | 7.8 | 8.8 | 8.9 |
| $n$ | 6.9 | 5.49 | 5.28 | 5.65 | 8.24 | 8.54 | 9.13 | 8.21 | 8.6 | 8.6 | 7.3 | 7.6 |

The effect of cholesterol is clearly indicated in Fig. 2 A, B, D, and E. Fig. 2 C and F show that mutant nematodes (without NMDA and GABA receptors) can also be anesthetized (although both strains at higher pressures). The number of body bends performed by the nematodes is also an important parameter to evaluate their motility [33]. Figures 3A and B show the body bends of CE as a function of gas pressure. Clearly, the body bends drop to zero indicating full anesthesia. Let us remark that during the anesthetic process, the nematodes get different bending conditions or contortions (Figs. 3C-F), eventually acquiring a linear conformation, see Fig. 3F. Related to the body bends, there is another parameter employed in nematode studies [33]: the number of reversals per minute (Fig. 4), which accounts for the number of times the animal changes movement direction. This correlates with the number of body bends. Indeed, reversals per minute also drop to zero, see Fig. 4. Finally, in Fig. 5 we plot the beat heart rates of DM for Xe and N_2_O obtained directly from heart images obtained by a microscope using a fast camera. This experiment, as far as we know the first one performed on an animal during anesthesia, gives us important information: 1) the beat rate steadily reduces from around 240 to around 160 beats per minute with both gases, 2) arrhythmias are produced by N_2_O (see movie S3), and 3) we measured the effect of the gases in the hearth area, see *SI Appendix*, Fig. S4.

**Fig. 3.**
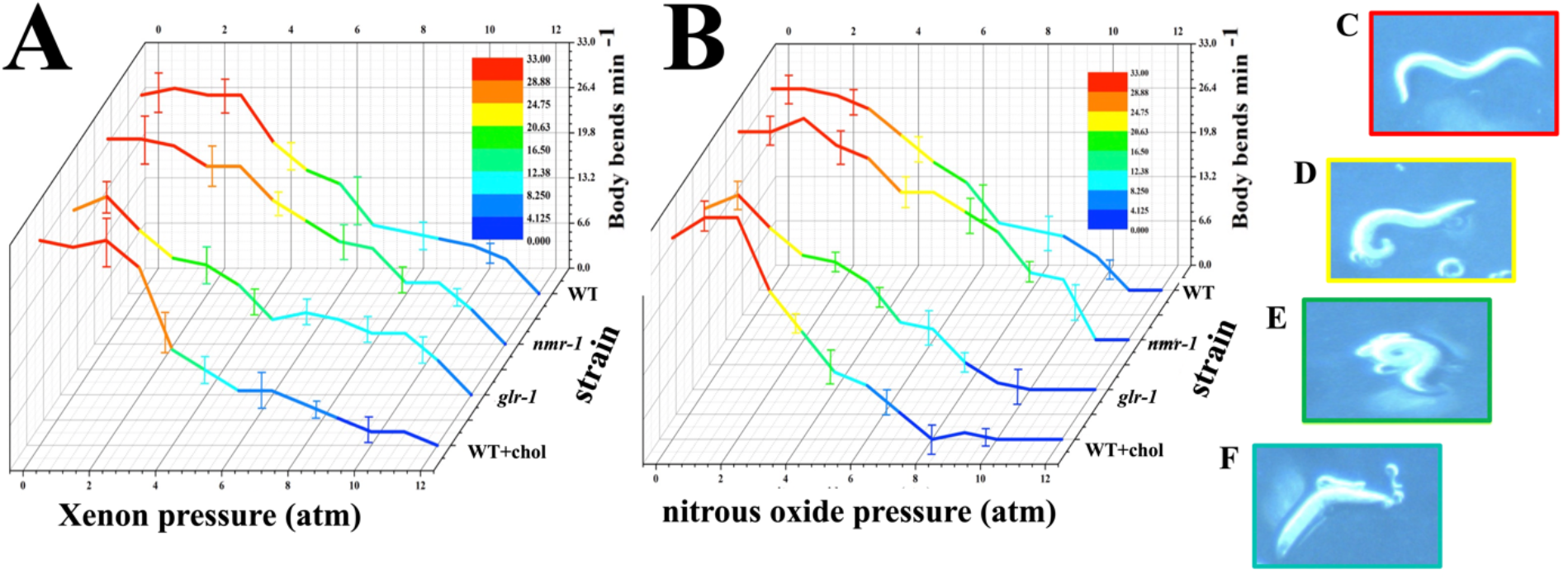
Body bends vs gas pressure for the different strains with Xe (A). Body bends vs gas pressure for the different strains with N_2_0 (B). Different conformations of the wild type of CE during anesthesia (C-F).

**Fig. 4.**
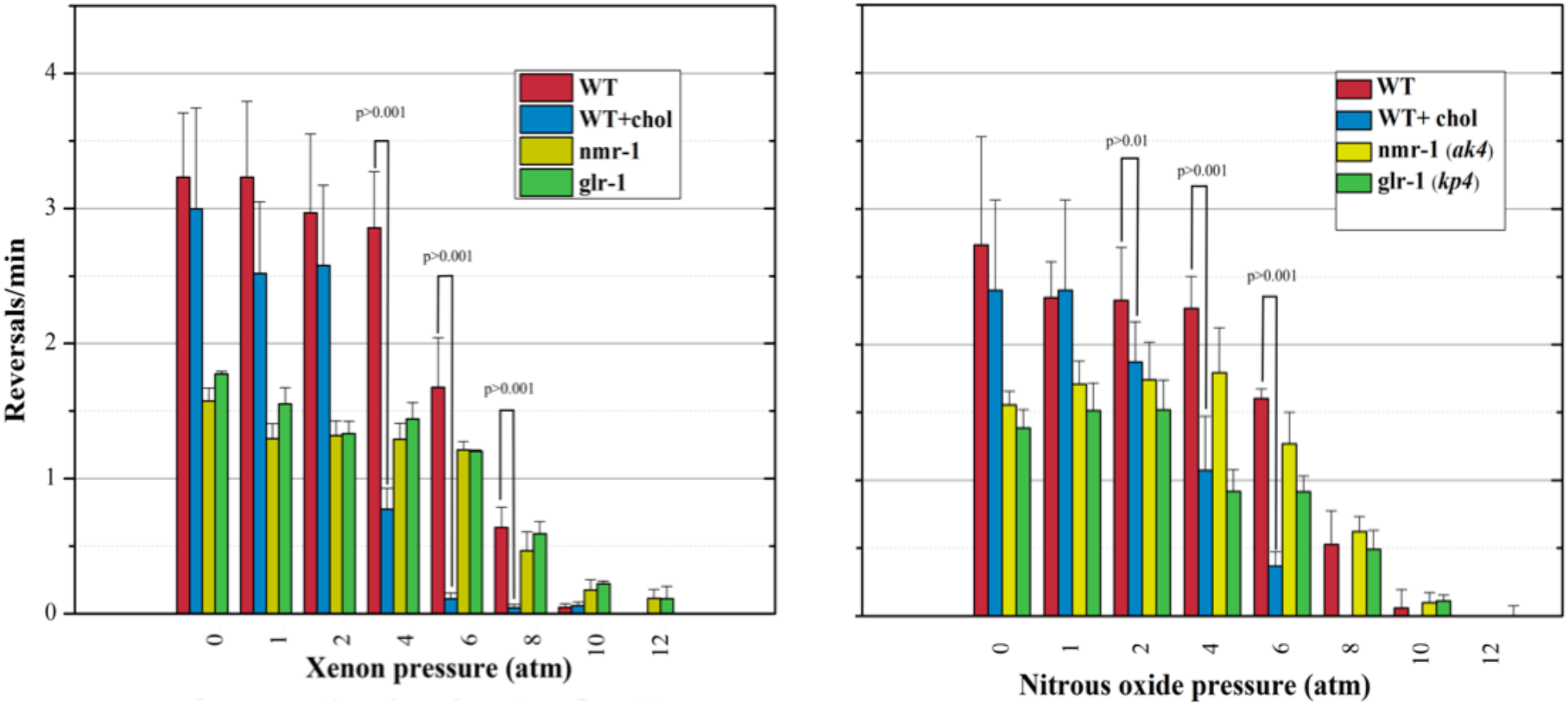
Reversals per minute for the different strains of CE with Xe (A). Reversals per minute for the different strains of CE with N_2_0 (B).

**Fig. 5.**
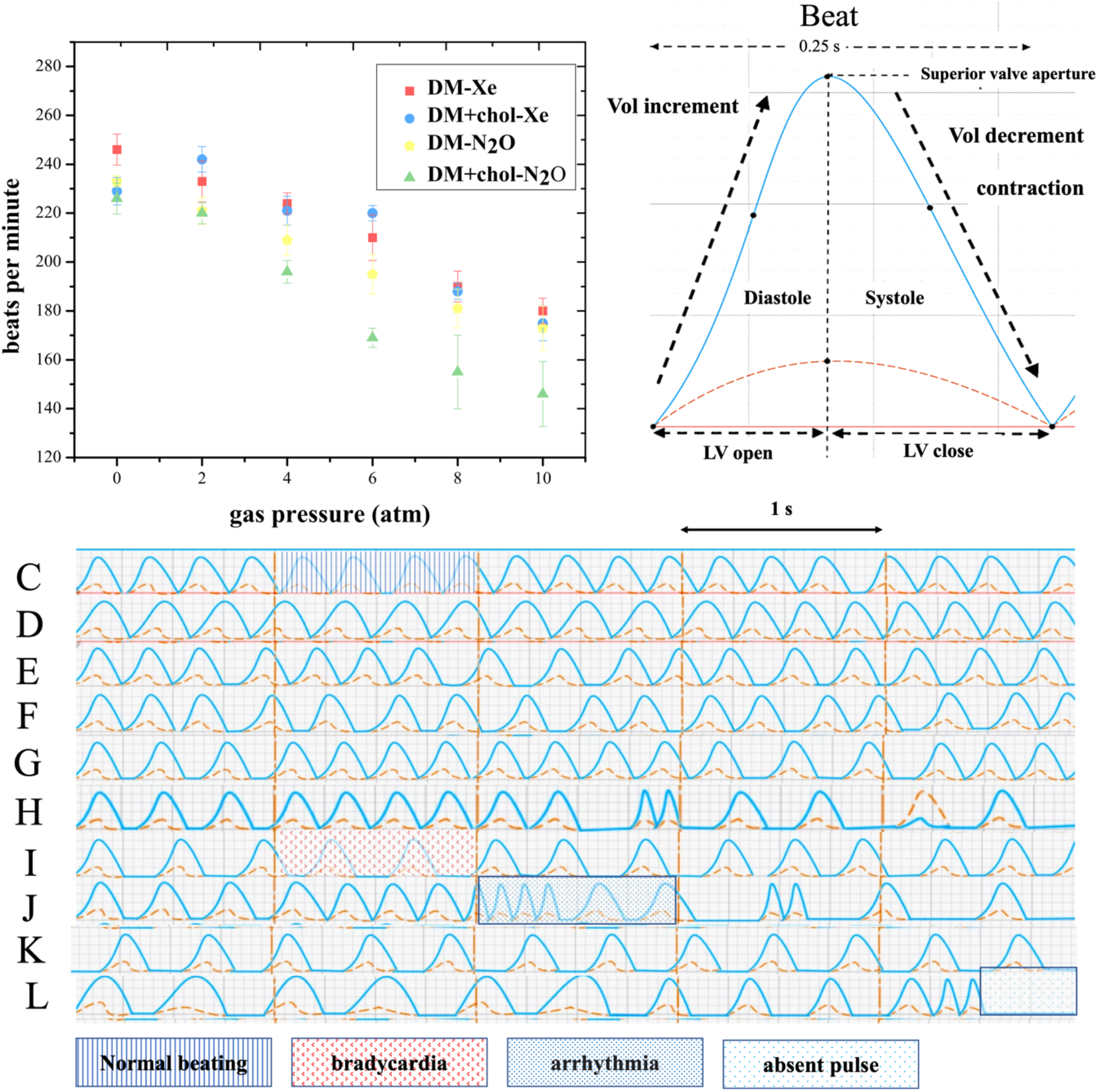
(A) Beats per minute of two animal groups (normal and extra cholesterol) after exposition to two anesthetic gases: xenon and nitrous oxide. (B) A beat starts with the aperture of the lateral valve, allowing hemolymph influx. Here cardiomyocytes are relaxed, and maximal cardiac area is reached (diastole). Systole starts once heart reaches its maximum volume (the lateral valve is closed), and the upper valve is opened to expulse hemolymph. A beat occurs in approximately 0.25s. A lapse of 5s was plotted for each group after its exposition to both gases during two stages of anesthesia: lack of half-motile response (EC_50_) and total immobility (EC_100_). Cardiograms of daphnids with a normal (C) and extra cholesterol diet (D), but no gases. (E and F) heart rate of the normal diet organism during EC_50_ for xenon and nitrous oxide, respectively. (G and H) heartbeat of DM feed with extra cholesterol after being exposed to Xe and N_2_O (EC_50_). (*I* and *J*) DM normal diet beating during total immobilization (EC_100_) induced by Xe and N_2_O correspondingly. (*K* and *L*) correspond to cardiograms of DM extra cholesterol diet during anesthesia produced by Xe and N_2_O.

## Discussion

We conducted experiments to test the anesthetic effect of xenon and nitrous oxide in the invertebrate *Daphnia magna* with a normal and enhanced cholesterol diet administered by a cholesterol nano-emulsion (see *SI Appendix*, Fig. S5). In order to perform the motility experiments, we modified the setup previously proposed and used in our group [34-35]. The method employs a technique to study in real-time, and under pressure, the organisms under study (*SI Appendix*, Fig. S1). We found that once the gases are in contact with the animals, they produce significant effects in the motor response. Krypton does not show an effect at the same used pressures, ruling out the impact of pressure in the immobilizing effect, see *SI Appendix*, Fig. S6.

The mechanism of *D. magna* to uptake gases in its aqueous medium is unknown. Nevertheless, the processes for blood oxygenation may explain how such gases exchange in the blood flow reaching the nervous system to induce immobility. Oxygen exchanges in *D. magna* occur near the posterior margin of carapace and in the part of the head [36]. Alterations produced by low oxygen levels were discarded due to the faculty of DM to adapt to a wide range of low-oxygen conditions [37,38]. Moreover, the experiment using krypton confirms this claim.

*Daphnia magna* presents a complex nervous system (see Fig. 1E), and not all the receptors have been reported, while the use of clones reduced the influence of genetic variations. We presumed that xenon and nitrous oxide would produce motor effects that resembled those observed using other volatile anesthetics [39]. Although most general anesthetics enhance the activity of GABAA (γ-aminobutyric acid type A) [40,41], the noble gas xenon interaction is unclear. It has been reported that xenon inhibits the excitatory NMDA receptor channels [42], but also this inert gas can inhibit GABAergic [43-45] and AMPA/KA receptors [46,47]. Lipid rafts are lipid microdomains rich in cholesterol and sphingomyelin, which are present in all cells. These sites are involved in protein trafficking and formation of cell signaling complexes and play an important role on G-protein mediated signal transduction and ligand-gated ion channels. In adult neurons, it has been reported that the NMDAr is distributed between postsynaptic density and lipid rafts [48]. However, these microdomains may contribute to the clustering of the GABAA receptor and the Na+, K+-ATPase at distinct functional locations on the cell surface [49].

The fact that cholesterol enhances GA (see Fig. 2) suggests that this physiological phenomenon is driven by lipid reorganization in the rafts rather than a selective inhibition of ion channels. Let us emphasize that cholesterol provided to the crustaceans and nematodes in the diet is easily transported to many parts of their bodies, including the nerve ring in CE [20]. Since we incubated both organisms for at least a week, cholesterol thus incorporates in the brain and nerve ring of the animals. Both species with the cholesterol diet were anesthetized with less pressure, see Fig. 2 A, B, D, and E. For example, the value of k for DM (CE) reduced from 7.02 (7.1) to 3.5 (4.9) atm for Xe. For nitrous oxide, the difference in k is much less pronounced, which can be explained by the fact that the interaction between N_2_O and cholesterol is lower. The logistic Hill’s equation (Eq. 1) has been extensively used to describe the relationship between drug effect (response) and drug concentration (dose). The Hill coefficient (n) estimates the number of ligands required to generate a response and k is the concentration to get a half-maximal effect. If n = 1, only one ligand or receptor is responsible for the response; n greater than one indicates multiple ligand binding [50] or unspecific action. The values obtained for n, for both gases, were greater than 5. In the insets of Fig. 2 (A, B, D, E) we depict how the dose-response curves would be if n = 1 (with the same value of k). Since the curves spread out, we employed a logarithmic scale. It is clear that n = 1 does not fit our data.

Since more cholesterol gives rise to more concentration of this lipid in the rafts[17-19], we argue that the enhanced GA is related to this increment. Pavel *et al*. found that a cholesterol depletion produced by methyl-β-cyclodextrin (MβCD) would have the same effect as a general anesthetic [3], while we found that more cholesterol enhances GA. Nevertheless, we claim our results do not contradict Pavel *et al*. findings: If more cholesterol exists, more anesthetic molecules arrive to the rafts and the enzymatic process to activate TREK-1 is enhanced. Furthermore, other channels like NMDA could also be affected as well by the increment of cholesterol in the lipid rafts. Indeed, since this lipid is highly hydrophobic, it could attract more xenon or nitrous oxide to destabilize the rafts through inward sequestration, as recently reported by us [9]. In other words, a higher amount of cholesterol first serves as a preferable zone of anesthetic nucleation and then, as Xe and N_2_O enter the rafts towards the tails zone, its sequestration or depletion occurs [9]. The effect of cholesterol is also clearly noted in the body bends and reversals per minute of CE, see Figs. 3 and 4.

Little is known about the effect of general anesthetics on the beat heart rates of animals. Here, we managed to observe the heart of the DM during the gas supply in real time. The beat rate drops from around 240 before anesthesia to 160 when the animals cease to move. The effect produced by Xe is different from the effect produced by nitrous oxide, which clearly produces heart arrhythmias. Campbell *et al*. saw hearth arrhythmias in DM during a lactose diet [51], presumably because lactose and other disaccharides directly affect sodium channels in its myogenic heart. From our results, we may conclude that nitrous oxide not only gives rise to GA but also affects ion channels in the heart of the crustaceans. But why N_2_O and not Xe? The answer is likely related to the number of hydrogen-bond acceptors (target recognition spots) of the lactose molecule, which has an elevated number (11), while N_2_O has 3 and Xe, 0. Interestingly, since Xe is an inert gas without the possibility of creating hydrogen bonds, we believe it does not selectively interact with ion channels, but only with lipid rafts where such channels are anchored. The fact N_2_O causes hearth arrhythmias in DM is indicative that this molecule not only acts on rafts, but also selectively with the ion channels.

Various important remarks are needed before closing. 1) Anesthesia was reversible in short times as soon the pressure is released; 2) Both species did not respond to krypton, therefore anesthetic effects are not produced by pressure, and 3) Since Xe is more hydrophobic than N_2_O, more pressure is needed to be dissolved in water during DM anesthesia (indeed, Henry’s proportionality constant, at room temperature, is 0,0043 mol/l bar, lower than 0,025 (mol/l bar) for N2O [52]). Yet, in terms of molarity, Xe anesthesia is more effective (24 mM) than N_2_O (110 mM).

Our results point out to a nonspecific action quantitatively described by a steep logistic behavior, which endorses the idea that general anesthesia cannot be driven by the selective inhibition of ion channels. Instead, GA is driven by the anesthetic influence of the physical properties of lipid rafts. Furthermore, when DM and CE are fed with cholesterol, anesthesia occurs at lower gas pressures, indicating its crucial role. Cholesterol is not a protein, therefore is not a target, explaining why Hill’s exponent is large. Finally, the strains *nmr-1* and *glr-1* can be anesthetized at higher pressures.

## Materials and Methods

### Gases

Xenon (investigation grade 5.5, CAS-No. 7440-63-3, analytical standard, purity > 99 %), nitrous oxide (CAS-No. 10024-97-2, clinical standard, purity > 99.5 %), and krypton (investigation grade 5, CAS-No. 7439-90-9, analytical standard, purity > 99 %) were acquired from Praxair Technology Inc (Mexico). All other chemical used were analytical grade and were purchased from Sigma-Aldrich (USA). Krypton was used as a control experiment.

### D. magna

The fleas were acquired from a permanent pond in Guadalajara, Mexico. They were acclimated and cultured in the laboratory for 3 weeks at 25 ºC under a 12-h light:12-h dark photoperiod and fed daily with unicellular culture *Chlorella vulgaris*. The cultured consist of a 1-liter glass bakers with 800 ml of a particular physiochemical characteristics medium (reconstituted water). This liquid medium was prepared with: 25 mM NaCl, 2 mM KCL, 1 mM MgCl2, 2 mM CaCl2, 2 mM MOPS as a pH buffer, and 1 mM NaHCO3.25 Medium pH and hardness CaCO3, was monitoring once a week (Thermo scientific Model Versastar Pro, USA). Dissolved Oxygen (DO) was determined in agreement to Mexican Official Norm, after 48 hours of aeration. Reconstituted water was refreshed twice a week and when necessary we replenished the lost volume by evaporation with MilliQ water.

### *C. elegans* wild type and strain culture

All worms were cultivated at 25ºC on nematode growth medium (NGM) which were previously seeded with *E. coli* OP50 bacteria. The wild-type strain used was N2 Bristol and was a gift from Rosa Navarro-González, Ph.D. professor at Physiology and Cell Development Department at the Universidad Nacional Autónoma de México, Mexico City. Transgenic strains VM487 [*nmr-1*(*ak4*)], KP4 [*glr-1(n2461*)], MC339 [*unc-64*(*md130*)], CB15 *[unc-8(e15) IV*], CB927 eDf28 (previously called *unc-24*(*e927*), were obtained from the Caenorhabditis Genetics Center (CGC, University of Minnesota, Minneapolis, MN, USA). VM487 is knockdown to *nmr-1*, a NMDA-type ionotropic glutamate receptor and KP4 is defective in an ionotropic glutamate receptor non-NMDA type.

### Drug application protocol

*D. Magna and C. elegans* were immobilized by volatile anesthetics in sealed chambers, see *SI Appendix*, Fig. S1. To discard effects produced by pressurization, a non-anesthetic gas (krypton) was used. Once we demonstrate pressure does not affect the motility, we administer both gases to hyperbaric conditions. To achieve hyperbaric conditions, we designed a special chamber (*SI Appendix*, Fig. S1) able to sustain up to 50 bars. The chamber allows us to observe, with a stereo microscope (10x, Axiovert-100; Zeiss, Gøttingen, Germany), the effect of the gases in real time.

### Image acquisition and processing

The captured films were transferred to a computer to create individual images. These were analyzed using the ImageJ software 1.46 with an autocorrelation plugin to compare the center of successive frames with the first frame and then calculate the autocorrelation ϕ (see Fig. 1).

Since at low pressures ϕ decays rapidly from 1 to 0 (the animals move normally) but barely decays at high pressures (they are anesthetized), we argue that the sum the autocorrelation values for all frames is a good quantitative measure to describe the motility state of the animals. Hence, to use the sum as a parameter constricted between 0 and 1 (needed to estimate the dose-response anaesthesia curves of the animals) the following heuristic equation was employed:

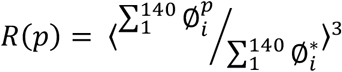

*p* stands for the pressure used in each experiment. Note that the data are normalized to the sum of the autocorrelation values for the highest pressure (10 bars), indicated by * (animals fully anesthetized). We used the power of 3 to strengthen the behavior between low and high pressures. The power could be any other number larger than 3.

### Cholesterol intake

Since cholesterol is a very hydrophobic molecule, it does not dissolve in the aqueous medium where the animals live. Therefore, we devised the following way to feed the animals with it. We dissolved a given amount of cholesterol (50 mg/ml) in laboratory-grade coconut oil. Then, we took 1 ml of the cholesterol/coconut stock and made a nanoemulsion with 100 ml of pure water (1:100). Using dynamic light scattering (DLS), we measured the size distribution of the oily nano drops, see *SI Appendix*, Fig. S1. The drops are small enough to be ingested by the animals.

### Statistics

Continuous variables are presented as means ± SD. Student’s t test and analysis of variance (ANOVA) tests were used for group comparisons of continuous variables as appropriate. For multiple comparisons, Bonferroni correction was used. Sigmoidal dose-response curves were fitted with Hill-Langmuir equation. Strength and direction of associations were performed through Spearman’s rank-order correlation. Linear regressions were assayed as appropriate. A two-tailed P value of <0.01 was considered statistically significant, and all statistical analyses were carried out with Origin Lab software (version 8.2).

## Supporting information

DM_motility_N2O

CE_motility_Xe

DM_heart_N2O

## ACKNOWLEDGMENTS

This work has been supported by grants FC-1132 (Conacyt) and FIDSC-100, Mexico. K.C.M.R.S. acknowledges a scholarship by Conacyt, Mexico. We thank Erel Levine for fruitful discussions. Fig. 1 D was created using BioRender.com.

## Notes

https://www.icloud.com/iclouddrive/035CxByeInOelVtJ0v5Q0zzUw#Supplementary

### Competing Interest Statement

The authors have declared no competing interest.

