## supporting information for "A high-cholesterol diet leads to faster induction of general anesthesia in two model animals: *D. magna and C. elegans*"

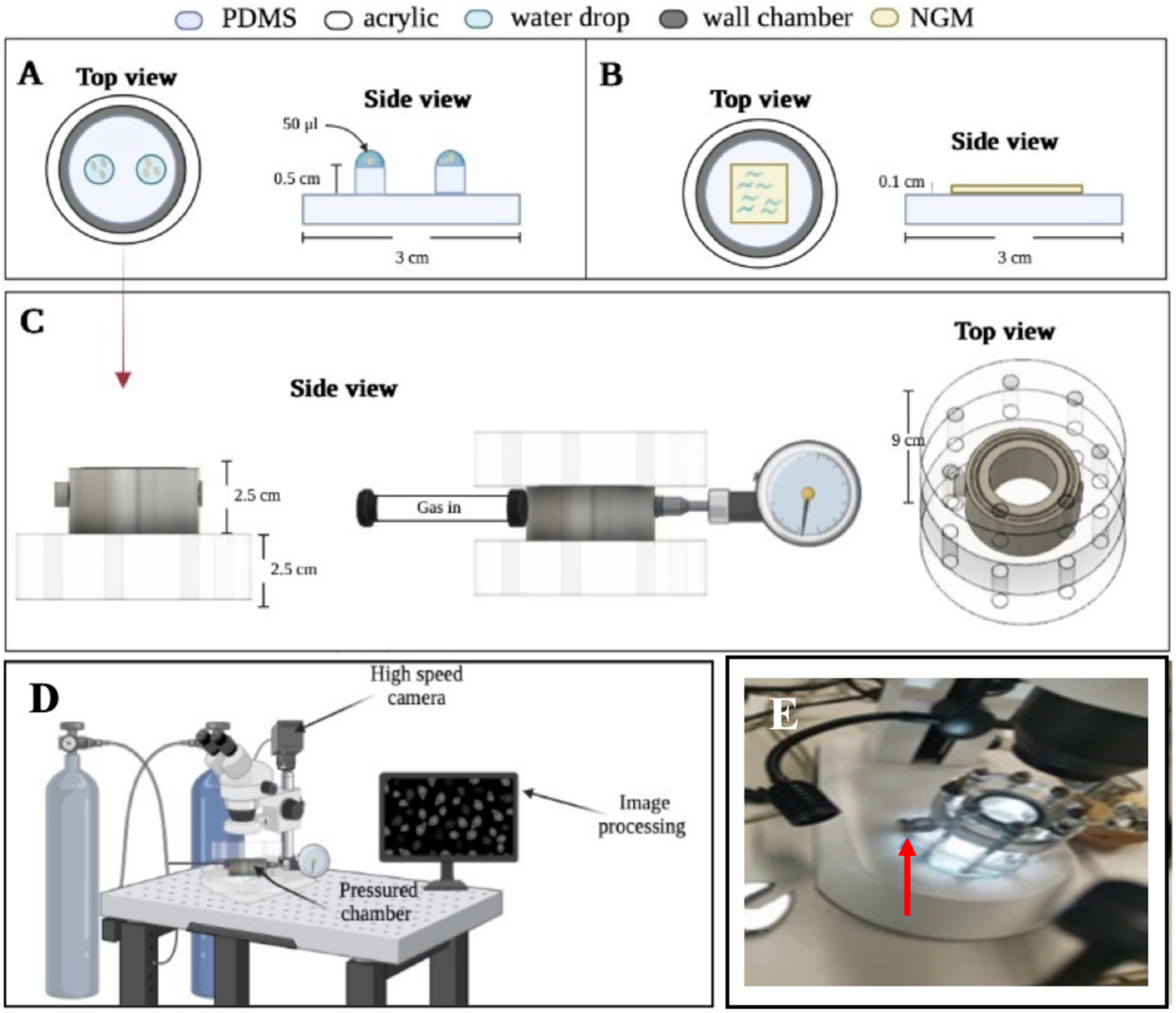


**Fig. S1** Experimental setup used in the experiments. (A) Top and side views of the post holding water drops with the fleas; (B) Top and side views of the surface with nematodes; (C) Side and top views of the pressure camera; (D) integrated setup, and (E) a photograph of the real cell mounted on the microscope.

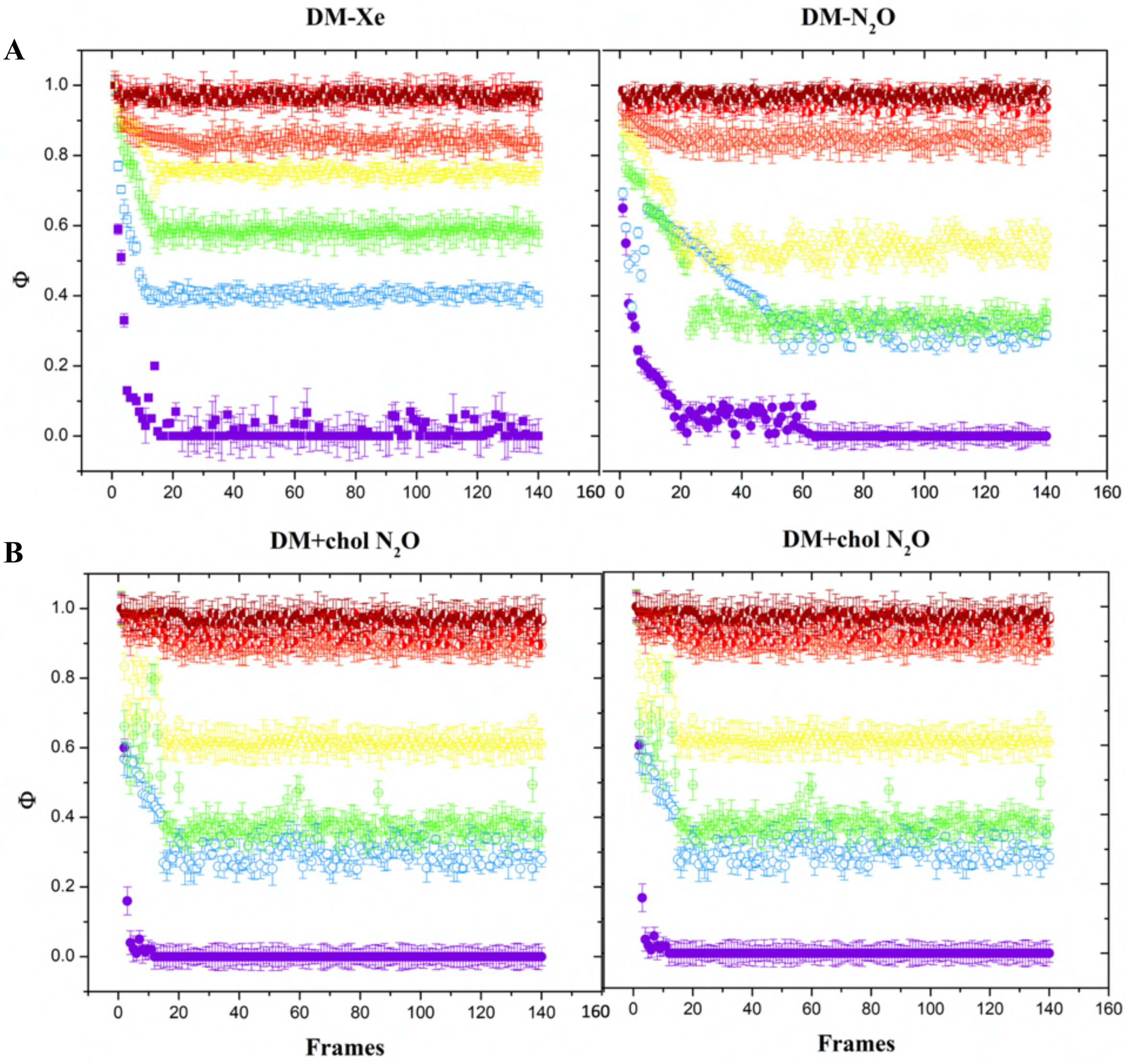


**Fig. S2** Autocorrelation values as a function of captured frame. Note that the values go from 1 to 0. A sudden drop from 1 to 0 (see for instance the experiments at 0 atm, purple dots) means that the animals moves normally. As the pressure increases, the autocorrelation values grow indicating that the daphnids get immobilized. (A) autocorrelation values on DM with a normal diet for xenon (left) and nitrous oxide (right). (B) shows the autocorrelation image values for the organisms feed with the cholesterol emulsion.

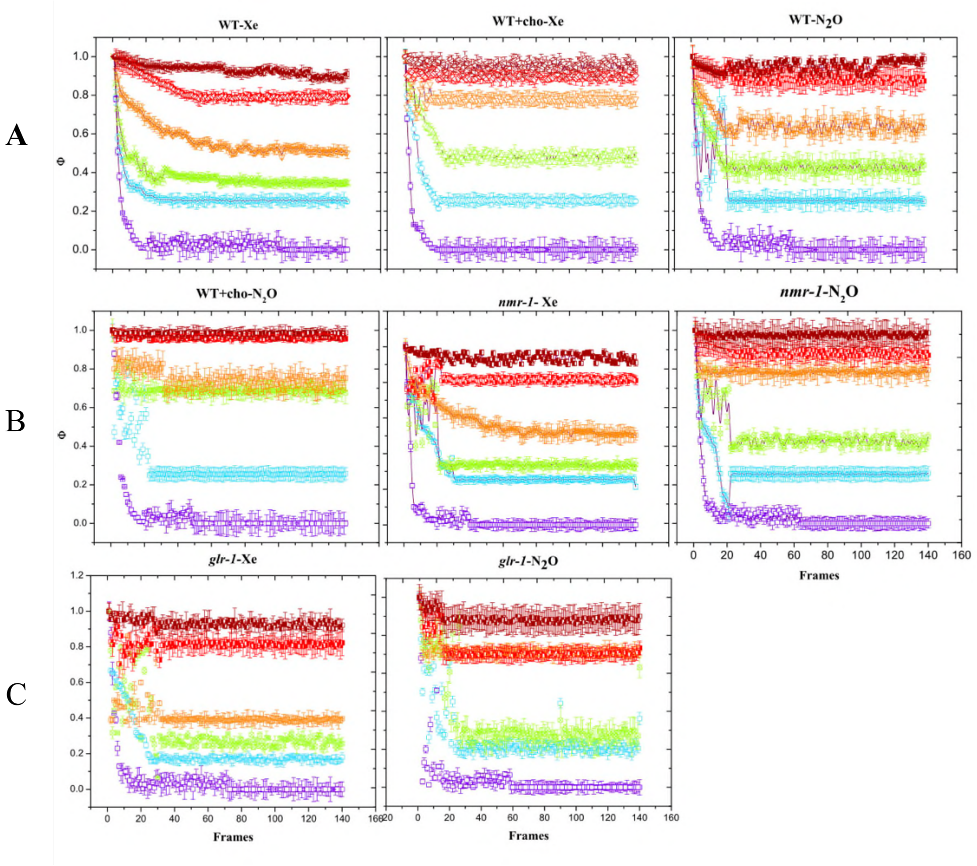


Fig. S3 Motility assay for *C. elegans* with nomal diet and feed with cholesterol. Each curve represents a different autocorrelation value(Φ): purple dots depict autocorrelation values in absence of gas, blue shows the values for 2 atm, green for 4 atm, orange for 6 atm, red for 8 atm, and wine for 10 atm of each gas. Autocorrelation assay for C. elegans show the motility with nomal diet and feed with cholesterol. Also, we depict the behavior for the strains vm487 and KP4.

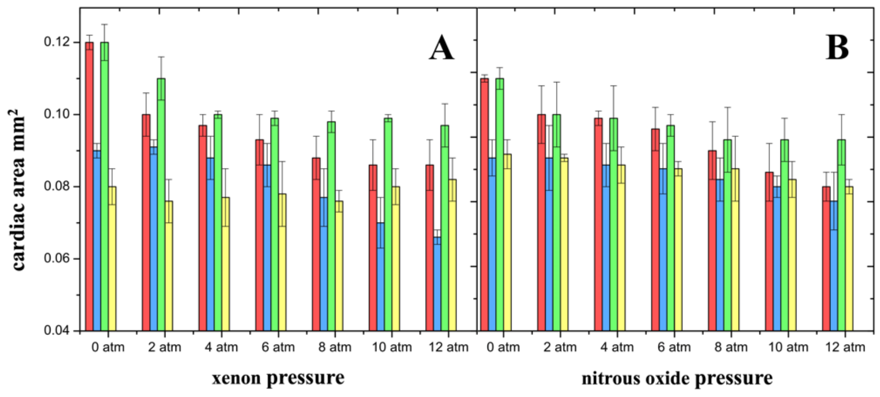


**Fig. S4** Cardiac area ($mm^2$) during cardiac cycle for two groups. DM normal diet Diastole (red) and systole (blue). For DM extra cholesterol diet diastole are in green bars and systole yellow ones. All groups were exposed to different pressures of the anesthetic gases xenon (left) and nitrous oxide (right.)

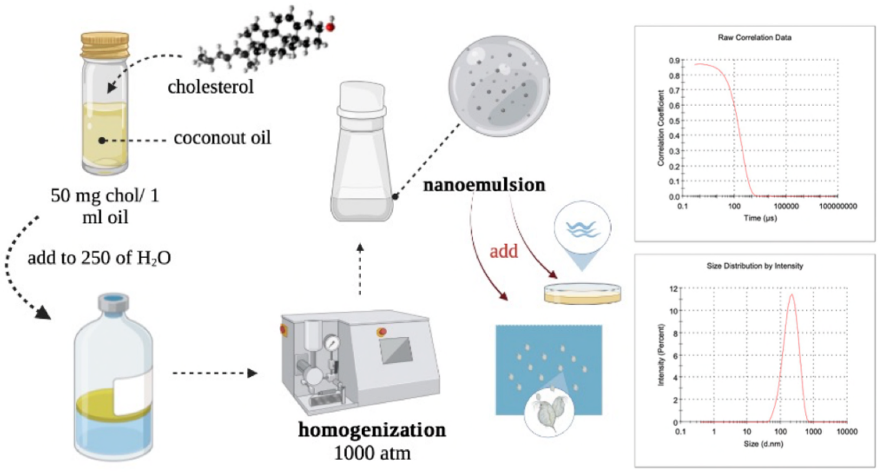


**Fig. S5** Preparation of the nanoemulsions. 250 mg were dissolved in 5 ml of coconut oil (Cocos nucifera 99%) CAS 8001-31-8. Then this mixture was added to 250 ml of milliQ water. Using a homogenizer operating to 1000 atm a stable emulsion was achieved. Size distribution of the nano-drops in water was measured centered at 250 nm. Two milliliters of the nanoemulsion were added directly in the Petri dish with wild type worms. 50 µl of the nanoemulsion was added to 50 ml of DM every day.

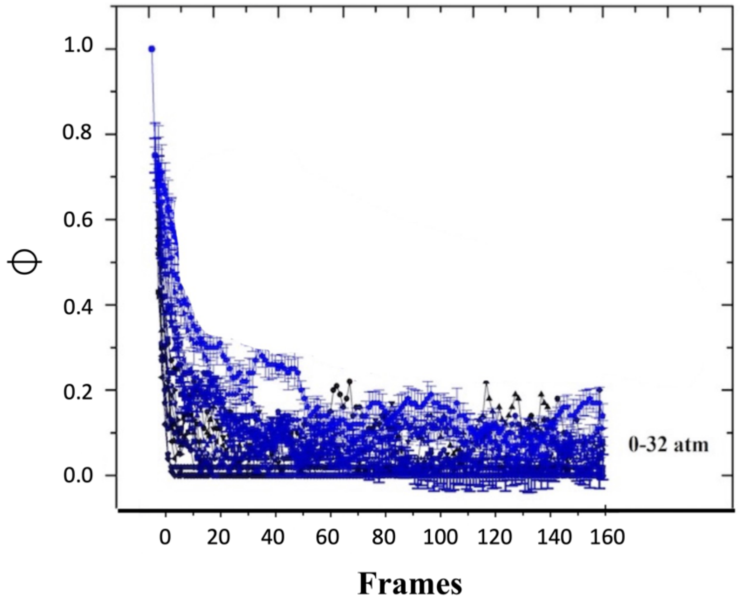


**Fig. S6** Autocorrelations values of DM for krypton, showing that this gas does not cause anesthesia at pressures in the studied range (0-12 atm). Even at higher pressures, the effect of krypton is negligible.
